# Limited Cell-Autonomous Anticancer Mechanisms in Long-Lived Bats

**DOI:** 10.1101/2024.02.29.582714

**Authors:** Fathima Athar, Zhizhong Zheng, Sebastien Riquier, Max Zacher, Dominic Alcock, Alex Galazyuk, Lisa Noelle Cooper, Tony Schountz, Lin-Fa Wang, Emma C. Teeling, Andrei Seluanov, Vera Gorbunova

## Abstract

Bats are remarkably long-lived for their size with many species living more than 20-40 years, suggesting that they possess efficient anti-aging and anti-cancer defenses. Here we investigated requirements for malignant transformation in primary bat fibroblasts in four bat species - little brown bat (*Myotis lucifugus)*, big brown bat (*Eptesicus fuscus*), cave nectar bat (*Eonycteris spelaea*) and Jamaican fruit bat (*Artibeus jamaicensis*) – spanning the bat evolutionary tree and including the longest-lived genera. We show that bat fibroblasts do not undergo replicative senescence and express active telomerase. Bat cells displayed attenuated stress induced premature senescence with a dampened secretory phenotype. Unexpectedly, we discovered that bat cells could be readily transformed by only two oncogenic perturbations or “hits”: inactivation of either p53 or pRb and activation of oncogenic RASV12. This was surprising because other long-lived mammalian species require up to five hits for malignant transformation. Additionally, bat fibroblasts exhibited increased p53 and MDM2 transcript levels, and elevated p53-dependent apoptosis. The little brown bat showed a genomic duplication of the p53 gene. We hypothesize that bats evolved enhanced p53 activity through gene duplications and transcriptional upregulation as an additional anti-cancer strategy, similar to elephants. In summary, active telomerase and the small number of oncogenic hits sufficient to malignantly transform bat cells suggest that *in vivo* bats rely heavily on non-cell autonomous mechanisms of tumor suppression.

## INTRODUCTION

Bats exhibit diverse lifespans [1, 2] with many species showing exceptional longevity. Longevity quotient (LQ) is the ratio of observed maximum longevity to the predicted lifespan normalized to the body size. Across Chiroptera, 65 species have a mean LQ of 3.52, which means that these bats live 3.5 times longer than an average non-bat mammal [3]; making them excellent models to investigate adaptations for longevity.

Cancer is a multistage process involving accumulation of genetic and epigenetic mutations in mitotic cells and the frequency of tumor formation depends on the number of cells divisions over time. Therefore, longer lifespans with more cell divisions, and longer exposure to exo- and endogenous stressors increase cancer incidence [4, 5]. However, despite their exceptional lifespans, few to no tumors have been reported in long-lived wild and captive populations of bats [6–10]. Several comparative studies have uncovered adaptations in long-lived species that have anti-cancer functions. These include regulating telomerase activity in large species [11–13], increase in tumor suppressor gene copies in elephants [14–16], decreased in non-LTR retrotransposition [17], transposon-triggered innate immune responses for cell clearance [18], enhanced DNA repair in long-lived rodents [19], and regulation of uncontrolled proliferation by unique mechanisms such as early contact inhibition in the naked mole rat, and massive necrotic cell death in the blind mole rat [20, 21].

Bat genomic studies revealed multiple adaptive changes in their immune systems [22–29]. Many of these changes temper inflammation (reviewed in [30]), which may have an anticancer effect. Additionally, signatures of positive selection were detected in tumor suppressors [31], DNA damage checkpoint and DNA repair pathway genes [22, 31], and growth hormones [32]. However, with the exception of a recently published study [33], malignant transformation has not been experimentally investigated in bat primary cells.

The number of oncogenic hits required for malignant transformation varies by species. Human fibroblasts require 5 mutational hits for malignant transformation, while mouse cells require only 2: inactivation of either Rb or p53 tumor suppressors and activation of Ras signaling pathway [34]. We previously investigated the numbers of oncogenic hits required for transformation of multiple rodent species. The number of hits was shown to increase with animal size and lifespan (ranging between 2 in the mouse and 5 in the beaver) reflecting stricter control over cell proliferation in longer-lived and larger-sized species [12].

In the present study, we sought to investigate anti-cancer mechanisms in bats. We isolated primary wing fibroblasts from four species of bats; little brown bat (*Myotis lucifugus*) with maximum lifespan (MLS) of 34 years, big brown bat (*Eptesicus fuscus*; MLS of 19 years), cave nectar bat (*Eonycteris spelaea*; MLS of over 8 years; related species *Eidolon helvum* have MLS of 22 years), and Jamaican fruit bat (*Artibeus jamaicensis*; MLS of 19 years). These species are from four bat families, span both bat suborders, and include the longest-lived genera *Myotis* [35]. *M. lucifugus* and *E. fuscus* are from the family Vespertilionidae, and *A. jamaicensis* is from Phyllostomidae, both lineages are within the suborder Yangochiroptera. *E. spelaea*, from the family Pteropodidae, is placed within the other suborder Yinpterochiroptera. We investigated telomerase activity, number of ‘hits’ required for malignant transformation, and response to stress-induced premature senescence (SIPS) in bat wing fibroblasts, alongside skin fibroblasts from mice and humans for comparison. We show that, surprisingly, bat fibroblasts require only 2 oncogenic hits for malignant transformation which include inactivation of either p53 or Rb pathways and activation of RAS. Furthermore, bat cells exhibit elevated p53 activity and higher p53 transcript levels which resembles anti-tumor adaptations observed in elephants [14, 15].

## RESULTS

### Primary bat fibroblasts and tissues are positive for telomerase activity

Telomeres in mammals are long tracts of tandem hexameric TTAGGG nucleotide repeats that protect the ends of the chromosomes and usually require a specialized RNA-dependent DNA polymerase, telomerase, to be replicated. Telomerase is repressed in somatic cells of large mammals [11] but is reactivated during tumorigenesis. Hence, suppression of somatic telomerase activity represents a tumor suppressor mechanism that evolves with large body size [36, 37]. We used Telomere Repeat Amplification Protocol (TRAP), a PCR-based assay, to test for telomerase activity.

Cell extracts tested by TRAP assay showed that all four bat species possess telomerase activity in their wing fibroblasts. *M. lucifugus* and *E. fuscus* fibroblasts showed more robust telomerase activity when compared to *E. spelaea* and *A. jamaicensis* (Figure 1a). We also confirmed this by testing several tissues we obtained from two of the bat species. Extracts from lung, spleen, wing, liver, heart, kidney, and brain tissues from *E. fuscus* and *E. spelaea*, showed telomerase activity. Within each bat, post-mitotic tissues like the heart and kidney showed lower activity than the lung, liver, spleen, and wing tissues, with the spleen showing the highest activity. (Figure 1b). Telomerase activity in the wing tissue extract of *E. fuscus* was higher than in *E. spelaea,* similar to the observations with respective wing fibroblast cultures. Bat wing fibroblasts proliferated continuously in culture and did not show replicative senescence, which is consistent with the presence of active telomerase (Figure 1c). We tested telomerase activity in wing fibroblasts from *M. lucifugus* and *E. spelaea* differing by at least 20 population doublings (PDs) (Figure 1d). We found small to negligible reduction in telomerase activity in fibroblasts differing by 20 PDs from both *M. lucifugus* and *E. spelaea*, suggesting that the telomerase activity is sustained over several cell divisions.

**Figure 1:**
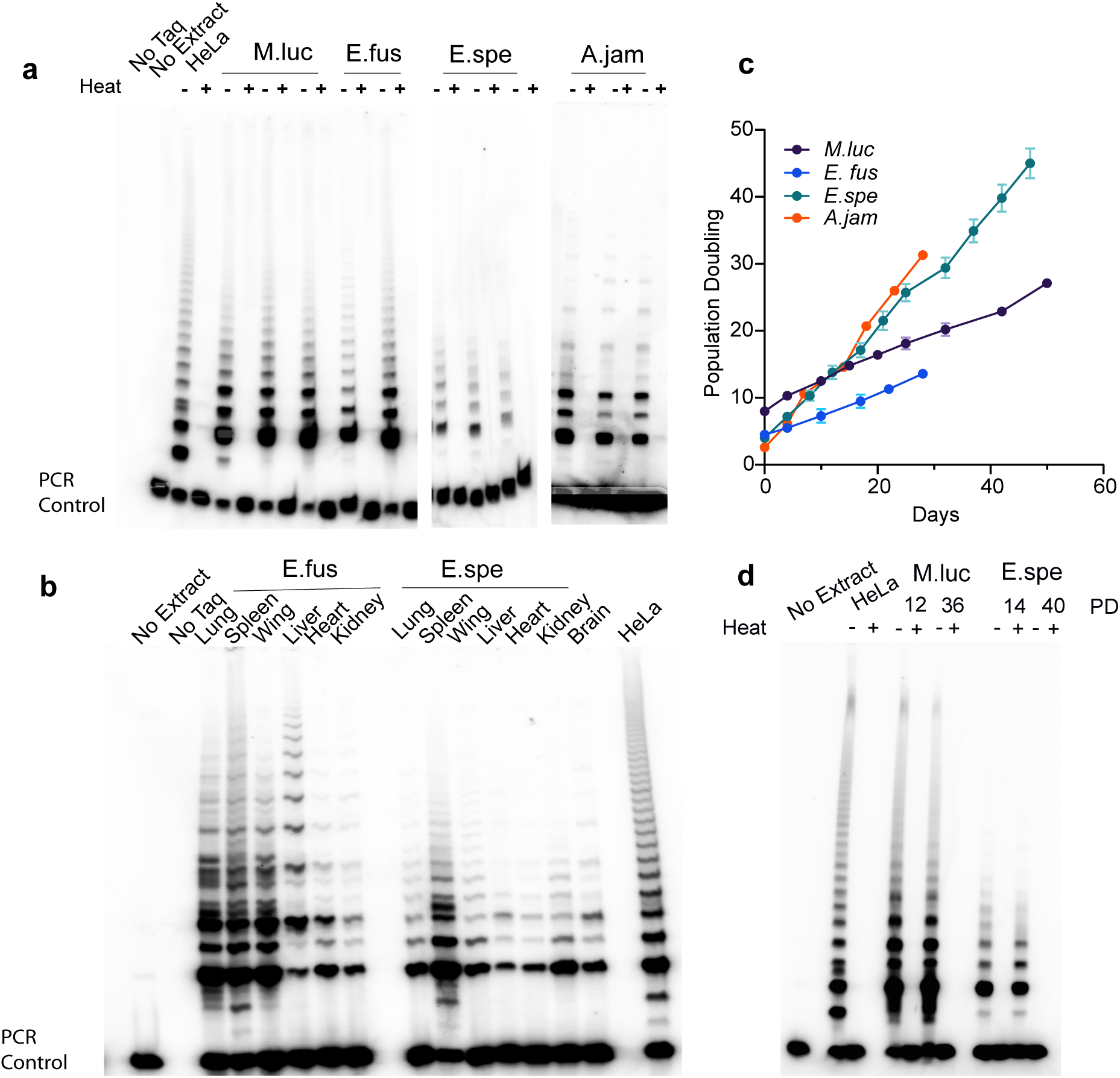
Bat cells and tissues possess telomerase activity. Telomerase Repeated Amplification Protocol (TRAP) assay is shown using wing fibroblast or tissue extracts from bat species listed. HeLa extract is used as positive control and heat inactivated extract of each sample is used as negative control. Internal PCR control is used to show absence of PCR inhibitors in samples. a, Telomerase activity in bat wing fibroblasts derived from four different species. b, Telomerase activity in several tissues of bats *E. fuscus* and *E. spelaea*. c, Representative population doubling curves for four species of bats showing fibroblast growth rates in culture. d, Telomerase activity in bat wing fibroblasts of *M. lucifugus* and *E. spelaea* differing by at least 20 population doublings.

A previous study assessed telomerase expression in blood and wing-punch-derived fibroblast transcriptomes. It hypothesized absence of TERT-mediated telomere maintenance in the long-lived bat *Myotis myotis* (MLS 37 years) given that relative telomere length did not change with age yet there was no evidence of significant TERT expression in the transcriptome data sets examined [1]. We tested wing fibroblast cultures from this population of *Myotis myotis* bats using the TRAP assay and found that fibroblast extracts were positive for telomerase activity (Supplementary Figure 1e). This suggests that telomere maintenance in *Myotis myotis* also follows canonical telomerase-mediated mechanisms with small bodied species expressing active telomerase, but without the expected increase in cancer incidence typical of other small bodied species.

### Bat fibroblasts require two oncogenic ‘hits’ for malignant transformation

Cell-type and species-specific differences have been demonstrated in the number of pathways to be perturbed (“hits”) for oncogenic transformation. The number and type of hits required for transformation may provide insights into species-specific inherent barriers to cancer formation *in vivo*. Human fibroblasts have been shown to require five to six pathways altered, while mice require two oncogenic hits for malignant transformation [34].

We sought to investigate if bat cells possess intrinsic mechanisms that confer resistance to malignant transformation. Wing fibroblasts from three bat species were used: *M. lucifugus*, *E. fuscus*, and *E. spelaea*. Stably transformed fibroblasts were generated by drug selection, expressing SV40 LT and its mutants - SV40-LT-K1 (inactivates p53 only) and SV40-LT-Δ 434-444 (inactivates Rb only), each in combination with HRas^G12V^. The combination of HRas^G12V^ with SV40 LT constitutes three hits, while HRas^G12V^ with either of the SV40 LT mutants constitute two oncogenic hits. Cells expressing GFP plasmid, SV40 LT-only, or HRas^G12V^-only were used as controls. Exogenous telomerase was not used because all these bat species showed intrinsic telomerase activity in our TRAP assays.

First, we analyzed anchorage-independent growth in soft agar assay. Surprisingly, overexpression of HRas^G12V^ along with either of the mutants of SV40-LT was sufficient for colony formation in all three bat species (Figure 2a). This suggests that two oncogenic hits - overexpression of Ras and inhibition of either p53 or pRb family of proteins - are sufficient for oncogenic transformation of bat fibroblasts. No colonies formed in controls.

**Figure 2:**
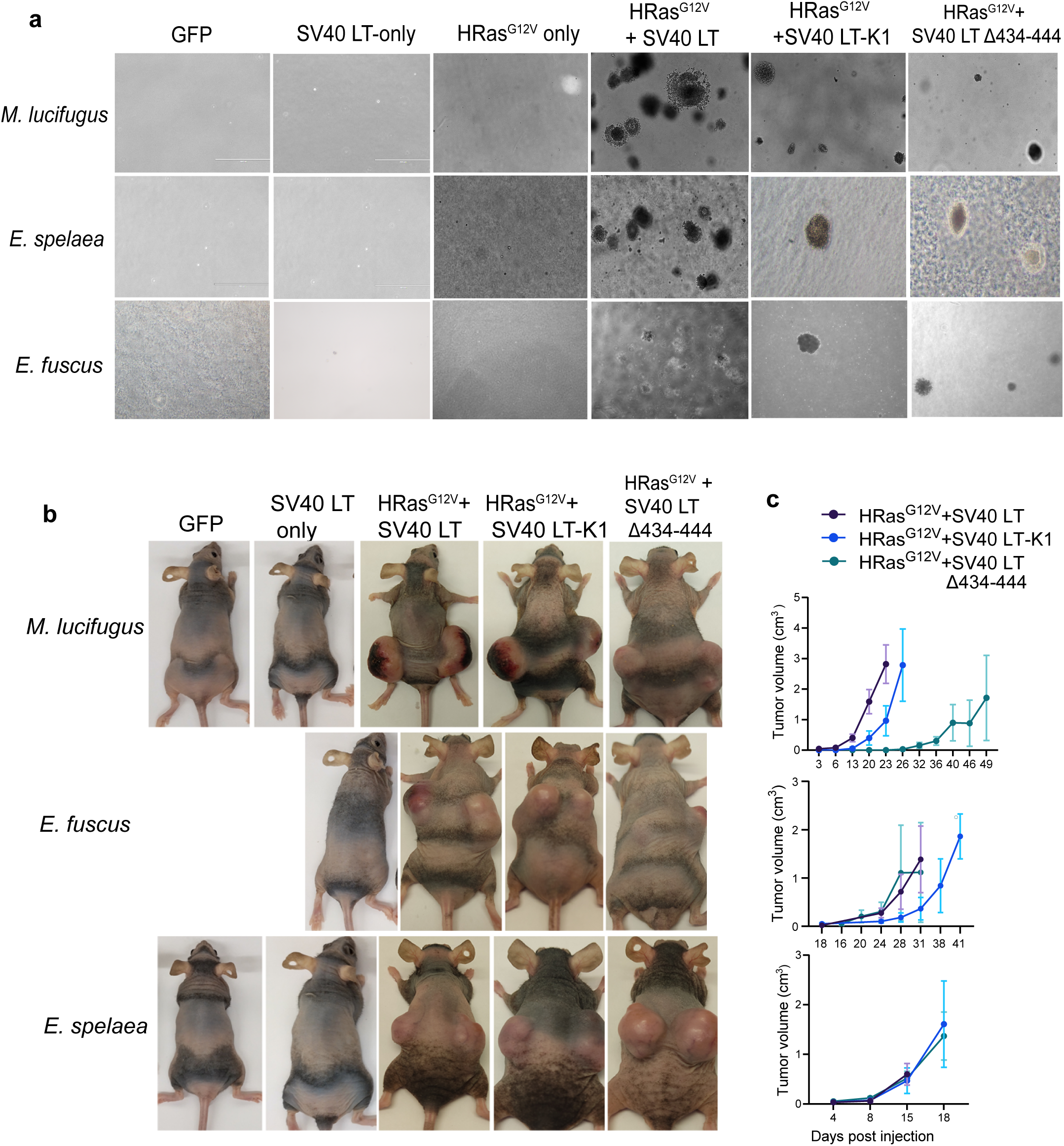
Two oncogenic hits are sufficient for transformation of bat fibroblasts. a, Representative images of colonies formed by bat wing fibroblasts from three bat species, *M. lucifugus*, *E. fuscus,* and *E. spelaea,* stably overexpressing oncogenes at the end of three weeks of soft agar assay. Scale bar, 400 μm. Two to three independent lines were tested for each species. Overexpression of HRas^G12V^ along with SV40 LT constitutes 3 oncogenic hits, while HRas^G12V^ along with either mutants of SV40 LT constitutes 2 oncogenic hits. b, Mouse xenograft assay using bat wing fibroblasts overexpressing oncogenes showing tumor formation in nude mice. One biological replicate per species was tested. Images shown were taken at the experiment end point. c, Tumor growth curves of transformed bat wing fibroblasts show increase in tumor volumes with time in mouse xenograft assay. Tumor volumes are an average of 8 injections performed for each cell line (4 mice/8 injections). Error bars show standard deviation (SD).

We further tested if transformed cells can form tumors in the mouse xenograft assay. No tumors formed in mice injected with control cells expressing SV40 LT or GFP only. Fibroblasts transformed with three forms of SV40 LT antigen along with HRas^G12V^ formed tumors in all three bat species, albeit with different tumor growth kinetics (Figure 2b). HRas^G12V^ with SV40 LT (three hits) grew the fastest (Figure 2c). Overexpression of HRas^G12V^ with either mutant of SV40-LT was sufficient for tumor formation. Thus, just as with laboratory mouse fibroblasts, a minimum of two hits were sufficient to transform bat fibroblasts.

### Bat fibroblasts display attenuated Stress-Induced Premature Senescence (SIPS) and instead undergo p53-dependent apoptosis

Stress-induced premature senescence, SIPS is the premature induction of senescence in cells using exogenous stressors. Two doses of γ-radiation (10 and 20 Gy) were used to induce SIPs in wing fibroblasts from four species of bats, dermal fibroblasts from humans, and laboratory and wild-caught mice, and the outcomes were compared. Cell proliferation was measured by BrdU incorporation assay three days post-radiation. Cells from all species underwent cell cycle arrest, with more than 50% reduction in BrdU positive cells at 10 and 20 Gy (Figure 3a). Day 12 post-radiation the cells were analyzed for induction of senescence by staining for Senescence-associated β-galactosidase (SA-β-gal). At 10 Gy about 40-60% senescent cells were observed in human and mice fibroblast cultures. However, the number of senescent cells in *M. lucifugus* and *A. jamaicensis* were four-fold lower (Figure 3b, c) compared to human cells. We found that the number of senescent cells in *E. fuscus* was higher and similar to mouse at 10 Gy. No interspecies differences in SA-β-gal were observed at 20 Gy where most cells of all species showed positive staining. *E. spelaea* was an exception, as it did not show positive SA-β-gal staining at pH-6.0 and therefore could not be quantified (Figure 3b). This could be because of species-specific differences in SA-β-gal. However, these fibroblasts showed a flat and enlarged appearance typical of senescent cells. In summary, some bat fibroblasts displayed attenuated levels of SIPS compared to mouse and human cells.

**Figure 3:**
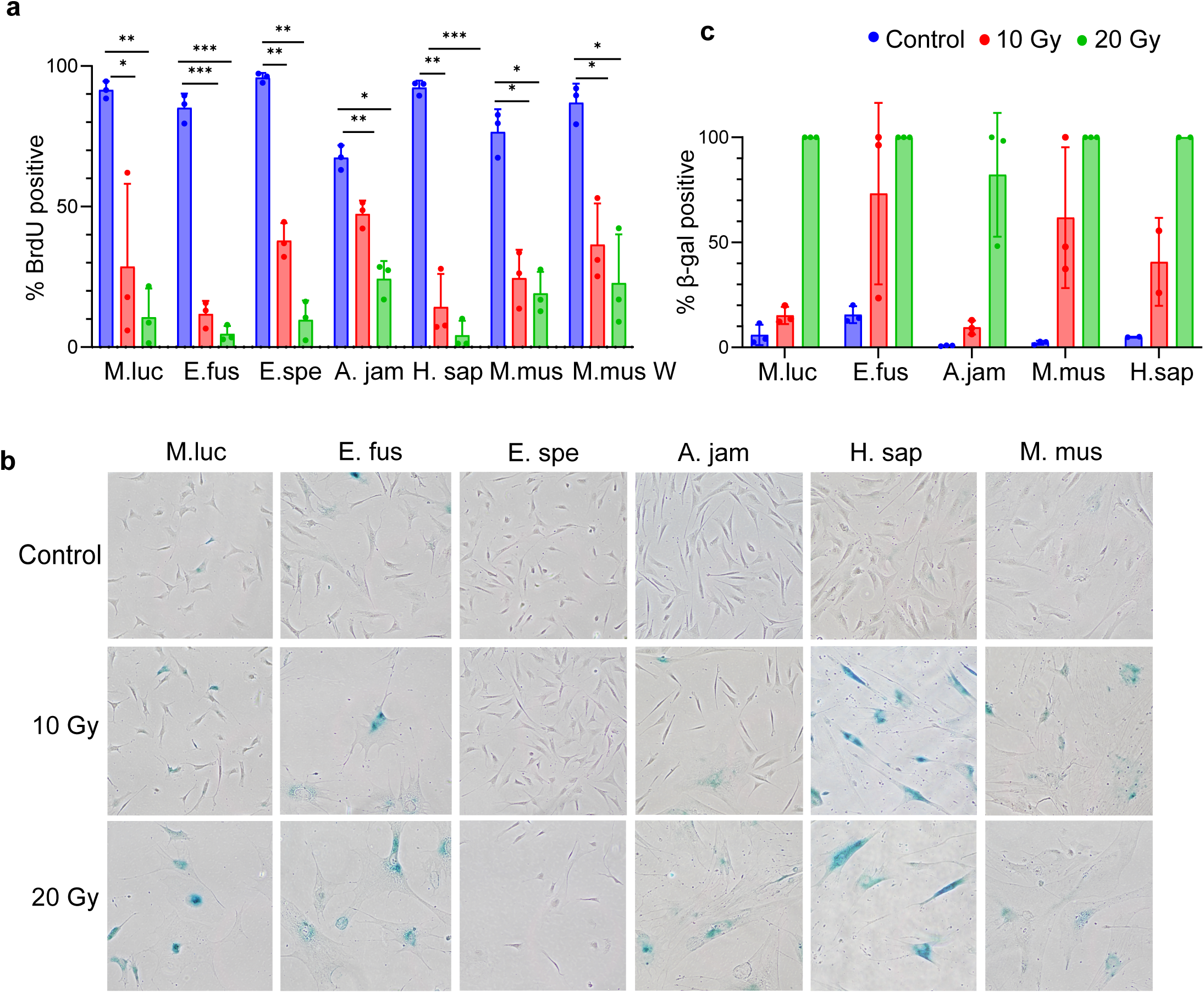
Analysis of senescence in bat species day 12 post-radiation. a, BrdU incorporation assay day 3 post-radiation in bat wing fibroblasts. n=3 for each species, error bars represent SD. \**p* <0.05, \*\**p*<0.01, \*\*\**p*<0.001 (Paired two-tailed *t*-test) b, Representative SA-β gal staining images of fibroblasts from all species 12 days post-radiation treatment. Bright field images taken at 10 x magnification c, Quantification of SA-β gal staining showing percentage of positively stained cells. n=3, shown as individual data points on plot. Error bars represent SD.

Senescent cells are characterized by the release of autocrine and paracrine secretory factors known as Senescence Associated Secretory Proteins (SASP) [38, 39]. Transcriptomes from fibroblasts treated with γ-radiation and allowed to senesce for 12 d were analyzed for SASP factors and other signatures of senescence. Gene Set Enrichment Analysis (GSEA) was performed using SASP gene sets from published literature [40–42]. At 10 Gy, *M. lucifugus* showed downregulation of several SASP factors that are generally upregulated during senescence (Figure 4a). This was also reflected in SA-β galactosidase staining, where *M. lucifugus* showed a lower percentage of senescent cells at 10 Gy. Detailed analysis of SASP gene expression showed that senescent *M. lucifugus* cells at lower doses of radiation altered expression of only about 30% of SASP genes compared to about 70% altered in mouse (Figure 4b). *E. fuscus*, *E. spelaea*, and *A. jamaicensis* also showed lower fraction of genes upregulated in some SASP categories when compared to mice in a similar analysis (Supplementary Figure 2). Because interferons are one of the components of SASP, we directly compared interferon expression in radiation treated cells from two different time points, 24 h and 12 days, post-radiation. Analysis of interferon expression revealed that senescent bat fibroblasts showed remarkably low interferon responses compared to human and mouse fibroblasts (Figure 4c).

**Figure 4:**
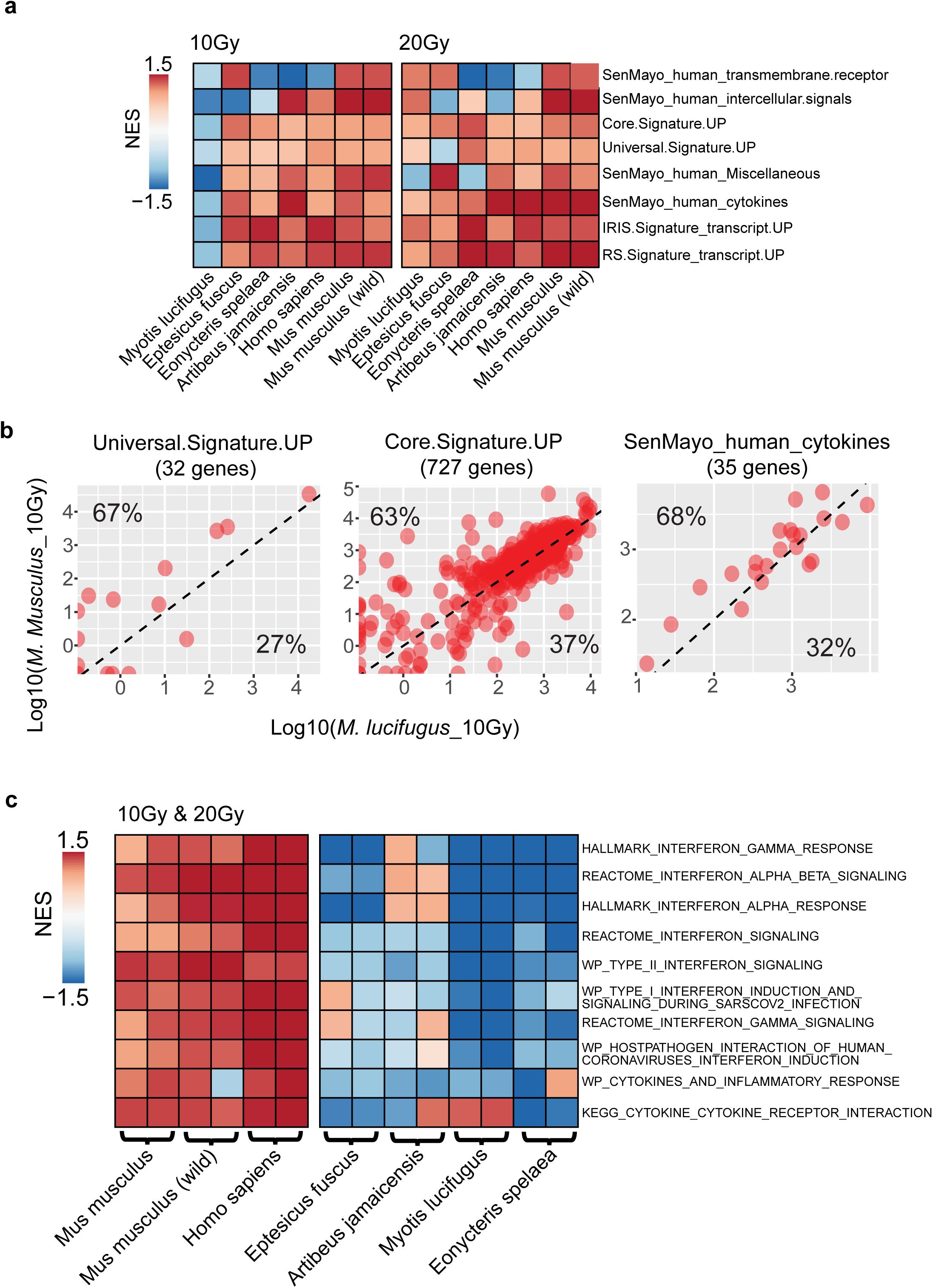
Comparative analysis of SASP in transcriptome of senescent fibroblast from bats, humans, and mice. a, Gene Set Enrichment Analysis (GSEA) of transcriptome using senescence-related gene sets day 12 post-radiation. Gene sets were collected from MsigDB database and published literature (See Methods). b, Scatter plots showing the expression differences between *M. lucifugus* and mouse senescent fibroblasts day 12 post radiation in 3 SASP-related gene sets. Percentages in the left-upper and lower-right in the plot indicate the fraction of genes with higher expression in mouse and *M. lucifugu*s, respectively. c, GSEA analysis of inflammation-related gene sets in senescent fibroblasts day 12 post-radiation. For this analysis, radiation-treated cells from 12 days were compared to those from 24 h post-radiation.

### Bat cells display elevated p53 activity

Genomic instability caused by γ-radiation treatment may lead to apoptosis. Apoptosis assays on day 3 post-radiation showed that most species of bats displayed significant increase in apoptosis upon irradiation, in contrast to mice and human cells (Figure 5a). Since we observed elevated apoptosis in bat cells following γ-radiation we hypothesized that bats may display elevated p53 activity. We tested p53 reporter activity in untreated fibroblasts from all species. We found that basal p53 transcriptional activity was higher in bat fibroblasts when compared to humans and mice. The vespertilionid bats, *M. lucifugus* and *E. fuscus* showed at least four-fold higher p53-reporter activity, while *E. spelaea* and *A. jamaicensis* showed about 1.5-2-fold higher activity compared to humans and mice (Figure 5b). Cells transfected with SV40-LT-K1 mutant, which specifically inhibits p53 function, demonstrated the specificity of the reporter activity. We then treated cells with Nutlin-3, an Mdm2 agonist, which stabilizes p53 by inhibiting Mdm2-mediated p53 degradation. Bat fibroblasts showed higher apoptosis than mouse and human fibroblasts following treatment with Nutlin-3 (Figure 5c).

**Figure 5:**
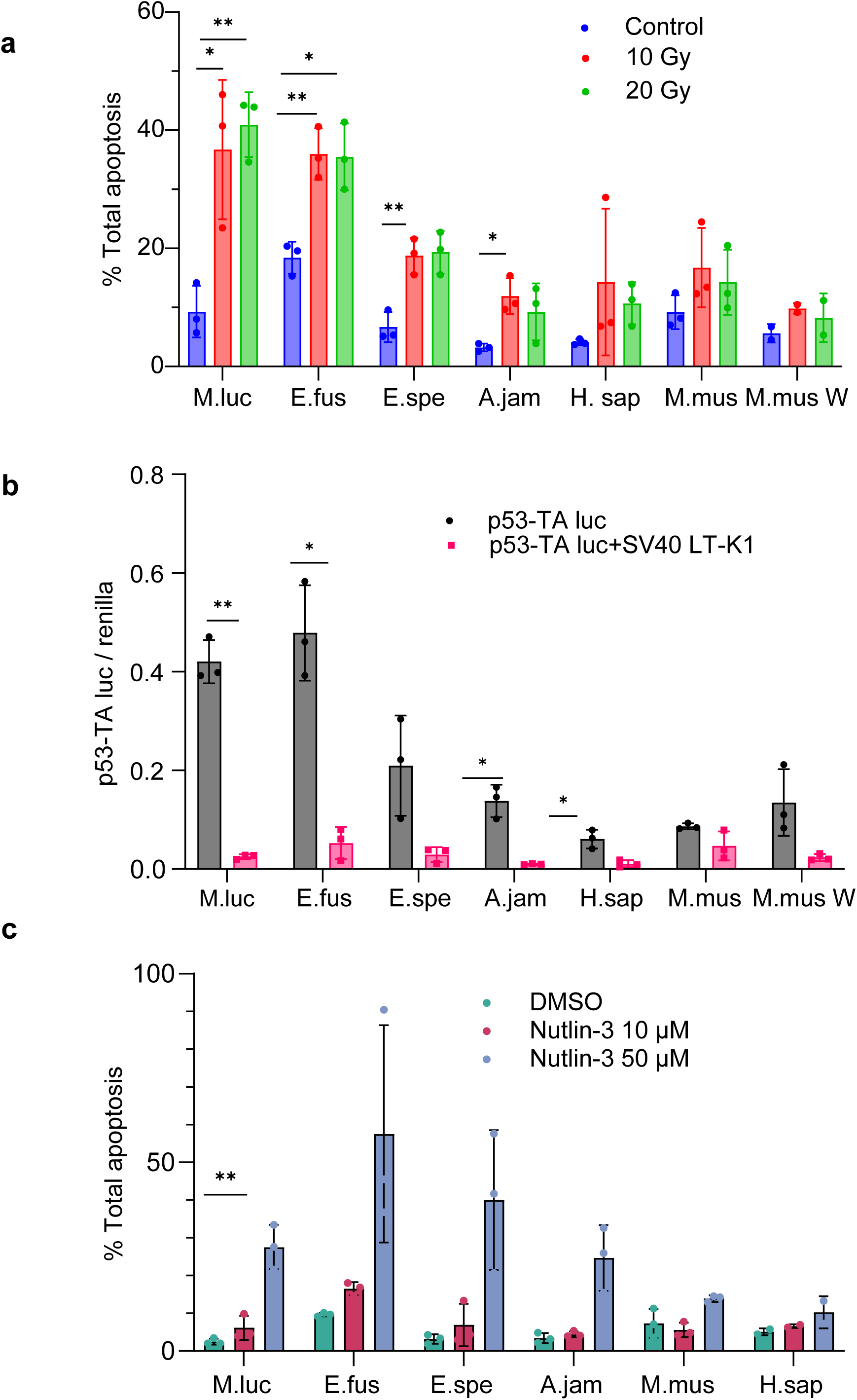
Bat fibroblasts undergo higher TP53-mediated apoptosis on treatment with γ-radiation. a, Annexin V apoptosis assay in bat fibroblasts treated with 10 and 20 Gy γ-radiation day 3 post-radiation treatment. b, p53-TA-luciferase reporter activity in normal proliferating bat wing fibroblasts and skin fibroblasts from mice and human. c, Annexin V apoptosis assay in proliferating cells treated with Nutlin-3 for 6 h. n=3 for each species, error bars represent SD. \**p* <0.05, \*\**p*<0.01 (Paired, two-tailed *t*-test)

We performed RNA sequencing of bat wing fibroblasts from *M. lucifugus*, *E. fuscus*, *E. spelaea*, and *A. jamaicensis* along with skin fibroblasts from laboratory mice, wild-caught *Mus musculus* mice and humans treated with 10 and 20 Gy doses of γ-radiation, at two time points, 24 h and 12 days post-radiation treatment. Both principal component analysis (PCA) and hierarchical clustering showed species-wise clustering of samples (Supplementary Figure 1a, b). Transcript levels of p53 were elevated in untreated bat fibroblasts compared to human skin fibroblasts (Figure 6a). The transcript profile of TP53 in skin fibroblasts from mice was comparable to that of bats, although they did not show higher transcriptional activity in the functional reporter assay in cells.

**Figure 6:**
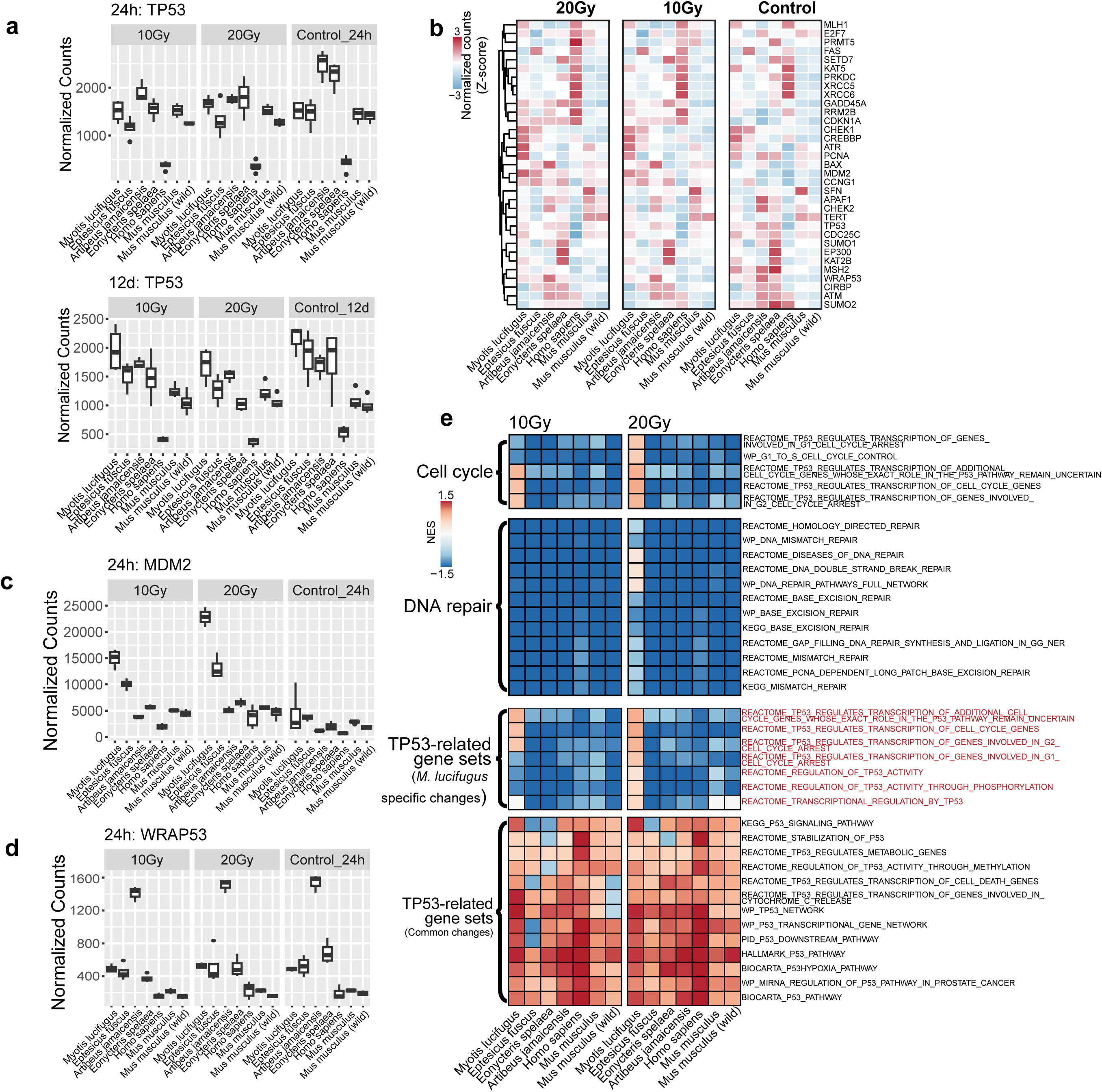
Functional enrichment analysis of TP53-related gene sets after irradiation. a, Boxplots showing *TP53* transcript levels in untreated control and radiation-treated cells at 24 h and 12 days post-radiation. The box plots display the median, the1st, and 3rd quartiles; the whiskers show a1.5×interquartile range. Data points outside the whiskers are outliers. b, Heatmap showing the expression changes of TP53 related genes 24 h after irradiation. c-d, Boxplots showing the expression changes of (c), MDM2 and (d) WRAP53 in each species at 24 h post-radiation. The box plots display the median, the1st, and 3rd quartiles; the whiskers show a1.5×interquartile range. Data points outside the whiskers are outliers. e, GSEA analysis for cell cycle, DNA repair, and TP-53 related gene sets in fibroblasts 24 h post-radiation.

We then analyzed TP53 transcriptional targets and regulators (Figure 6b). *M.lucifugus*, *E. fuscus*, and *A. jamaicensis* showed upregulation of *BAX*, a direct transcriptional target involved in p53-mediated apoptosis along with BCL2 [43]. *MDM2* (mouse double minute 2) is a key p53 antagonist that regulates p53 in a ubiquitination-dependent and independent manner [44, 45]. *MDM2* transcripts were upregulated in *M. lucifugus* and *E. fuscus* (Figure 6c). *WRAP53* (WD40 encoding RNA anti-sense to *TP53*) is a natural anti-sense transcript nested in the *TP53* gene [46, 47]. All bats showed higher levels of *WRAP53* transcripts when compared to humans and mice, in both treated and untreated cells. *A. jamaicensis* shows higher levels of *WRAP53* transcripts compared to other bat species (Figure 6d).

DNA repair genes have been shown to be positively selected in bats [22, 48]. Genes involved in DNA damage *(ATM, ATR, ChEK1, GADD45A, PCNA*) and cell cycle arrest genes (*CDKN1A, CCNG1*) were upregulated in most bat species (Figure 6b). PCNA showed higher upregulation in *M. lucifugus* in radiation-treated cells. Basal levels of Cold-induced RNA binding protein (CIRBP) with roles in DNA double-strand repair and stabilization of transcripts involved in cellular stress levels were higher in *E. fuscus, E. spelaea*, and *A. jamaicensis* compared to humans and mice [49] (Figure 6b).

We further performed GSEA using different p53 pathway gene sets. Since p53 is the universal pathway activated on radiation treatment, all species showed enrichment of p53 pathways terms (Figure 6e). However, *M. lucifugus* showed higher enrichment for some of the TP53-related pathway terms compared to other species. For example, genes of Reactome terms like “Transcription of cell cycle genes, G2, G1-cell cycle arrest”, “Regulation of TP53 activity by phosphorylation” were upregulated in *M. lucifugus*. The proapoptotic term “Reactome TP53 regulates transcription of genes involved in cytochrome C release” was also highly enriched in *M. lucifugus* at 10 Gy (Figure 6e).

GSEA for different pathways involved in DNA repair showed that *M. lucifugus* cells treated with 20 Gy showed slightly higher enrichment for DNA repair pathway terms at 24 h post-radiation treatment (Figure 6e). However, overall, the DNA repair pathway terms were not specifically enriched in all bat species in our data sets.

### Multiple copies of p53 in *M. lucifugus*

A p53-mediated apoptotic response to SIPs was also observed in a previous study with elephant fibroblasts. It was shown that the expansion of *TP53* copy number in elephants (20 copies) with several *TP53* retrogenes showing expression could be responsible for this phenotype [14, 15, 50]. We analyzed genomes of several bat species to see if *TP53* copy number expansion has occurred in Chiroptera (Figure 7a). We found that several bat species have 2-4 copies of *TP53*. Most intriguingly, the vespertilionid bats in this study, *M. lucifugus,* and *E. fuscus,* seem to possess 7 and 2 copies of *TP53*, respectively. Of the seven copies of *TP53* in *M. lucifugus*, one full copy duplication and five short retrocopies seem to exist in the available genome assembly, Myoluc2.0 (GCA_000147115.1). Potentially erroneous copy number estimation can result from a fragmented genome assembly. Therefore, we further analyzed and found the two copies in the HiC guided assembly of *M. lucifugus* genome (Myoluc2.0_HiC) [51, 52]. We additionally performed Cas9-targeted sequencing using guides directed to TP53 copies in *M. lucifugus* genome and obtained one long read (118 K) covering the *TP53* region of *M. lucifugus* in our preliminary analysis. This long read sequence suggests the presence of two full length copies of *TP53* (Figure 7b).

**Figure 7:**
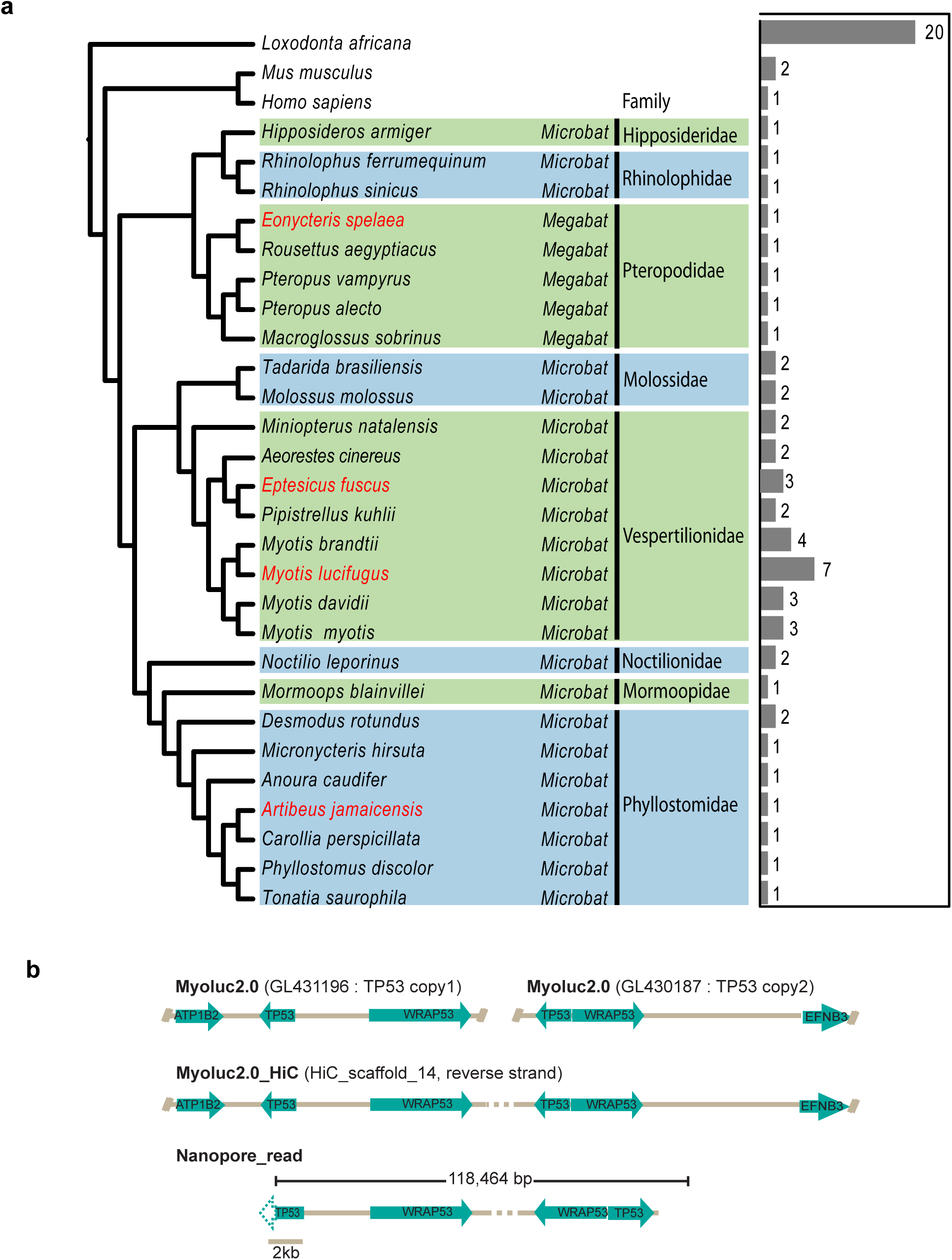
Analysis of *TP53* copy numbers in bats. a, TP53 copy numbers are shown as the bar at the right side. Phylogenetic relationship between species were collected from https://vertlife.org/. b, TP53 full copy duplications from genome, HiC-guided assembly and long-read sequencing in *M. lucifugus*.

## DISCUSSION

Bats are very long-lived for their size, and there are few to no reports of tumors in bats [6–10]. The number of oncogenic hits required for malignant transformation highlights species-specific inherent barriers to cancer development. For example, human fibroblasts have been shown to require five oncogenic hits for transformation (activation of telomerase, oncogenic Ras signaling, inactivation of p53, Rb and PP2A), whereas mice require only two (inactivation of either Rb or p53 and activation of Ras signaling) [34]. The number of oncogenic hits generally increases with body size and lifespan. For example, chinchilla and porcupine fibroblasts require 3 hits, naked mole rat fibroblasts require 4 hits, and beaver fibroblasts require 5 hits [12]. Here we discovered, unexpectedly, that all three species of bats tested, *M. lucifugus*, *E. fuscus*, and *E. spelaea,* representing the two bat suborders, underwent malignant transformation with two hits - the expression of HRas^G12V^ and inactivation of either Rb or p53. This suggested that bat fibroblasts are easily transformed and do not possess any peculiar inherent barriers to cell transformation. While we were drafting this manuscript, a similar study was published reporting that fibroblasts from seven different bat species, native to the Asian continent, undergo malignant transformation with overexpression of HRas^G12V^ and SV40LT antigen [33], indicating that this ease of malignant transformation is a wide spread phenomenon in bats. Transformed colonies of one of the seven bat species, *Myotis pilosus,* a member of the longest lived genera of bats, proliferated more slowly which led the authors to identify decrease in expression of *HIF5*, *COPS5*, and *RPS3* as potential mechanisms of cancer resistance in this particular bat [33]. However, this does not explain the paradox why species whose cells can be transformed with only two oncogenic hits show cancer resistance *in vivo*.

Downregulation of somatic telomerase activity is a tumor suppressor mechanism that evolved in mammals with the body mass larger than 5-10 kg [36]. Cancerous cells in these species must re-activate telomerase or use alternate telomerase pathways for survival [12, 53]. The presence or absence of telomerase activity in somatic cells and tissues has been tested only for a few bat species [13]. It has been observed that transcript levels of *TERT* are generally low in somatic cells, and PCR-based enzymatic assays are a better readout for telomerase activity. Having tested five bat species representing different clades, using the TRAP assay we confirm that bats have telomerase activity in their somatic cells and tissues, which is consistent with their body mass being below 5 kg. Within the five bat species that we tested, smaller bats *M. lucifugus, Myotis myotis,* and *E. fuscus,* (body weight 5-40 g) showed higher telomerase activity than larger bats, *E. spelaea,* and *A. jamaicensis* (40-80 g). This fit the general negative correlation between somatic telomerase activity and body mass, independent of lifespan [36].

Although bat cells expressed telomerase and did not experience replicative senescence, they could be induced to undergo SIPS using γ-radiation. Senescence is characterized by irreversible cell cycle arrest and release of pro-inflammatory factors called senescence-associated secretory phenotype (SASP) by the senescent cells [38, 39, 54]. The senescence program is a double-edged sword, though it prevents inheritance of mutations precluding potential malignant transformation, it also promotes age-related inflammation contributing to the aging process [55]. *M. lucifugus* showed distinct features in our analysis of SASP in senescent bat fibroblasts. Senescent *M. lucifugus* cells showed a markedly low SASP factor expression, suggesting that this bat may avoid the negative aspects of senescence. All bat species exhibited a blunted inflammatory response, with significantly lower interferon expression than non-bat species (Fig. 6d). Bats show reduced pro-inflammatory responses, contraction of type I interferon family and constitutive expression of interferon-α genes in cells and tissues, loss of PYHIN locus, and dampened activation of NLRP3 inflammasome [22, 30, 56, 57]. These unusual immune adaptations in bats, which may have evolved to counteract damage due to increased metabolism and higher body temperatures during flight, or for the co-existence with viruses may have been co-opted against the inflammation induced by senescent cells.

We observed that bat fibroblasts when exposed to genotoxic stress displayed enhanced apoptosis when compared to mice and humans. p53 is the primary responder to genotoxic stress and can induce apoptosis on extensive DNA damage [58]. We found that bat fibroblasts have enhanced basal p53 transcript levels. All four species had higher levels of p53 transcripts and showed two to four times higher basal transcriptional activity than in human or mouse cells. Higher transcript levels were sustained in cells upon radiation treatment and induction of senescence. Direct transcriptional targets of TP53 like *CDKN1A*, *CCNG1* (cell cycle arrest), *MDM2* (p53 regulator), and *BAX* (proapoptotic factor) [59] were upregulated in most species, albeit with enhanced levels in some bats.

How do bat cells tolerate higher p53 protein levels in their cells? Our transcriptome analysis provides few clues, although no universal pattern applicable for all four bat species emerges. Bat fibroblasts treated with radiation upregulated *MDM2*, with most upregulation seen in the vespertilionid bats *M. lucifugus* and *E. fuscus* (Fig. 5d). MDM2, a transcriptional target of TP53, is a E3 ubiquitin ligase that negatively regulates TP53 activity by ubiquitin-dependent and independent mechanisms [44, 45]. Interestingly, bat specific nucleotide changes in nuclear localization signal of TP53 and MDM2 are reported [22]. *WRAP53* is a natural anti-sense transcript of *TP53* and, in a non-reciprocal manner, positively regulates p53 activity by preventing degradation of *TP53* mRNA [46, 47]. All four bat species show enhanced basal levels of *WRAP53*, sustained following radiation treatment, and not observed in humans or mice. *A. jamaicensis* shows the highest expression of WRAP53 expression. Additionally, p53 was shown to undergo positive selection in bats, which may alter the pattern of transactivation by changing some of the numerous sites on p53 that undergo posttranslational modifications providing a fine tuning of p53 activity in response to various stimuli [22]. Interestingly, FBX031 gene which promotes degradation of MDM2 to increase p53 levels [60] underwent massive expansion in microbats with *Myotis lucifugus* and *Myotis brandtii* genomes containing over 50 copies [32, 61]. This expansion may also increase p53 levels and activity.

Enhanced reliance on apoptosis has been observed in the other long-lived and cancer-resistant species, the elephants. It was shown that the expansion of *TP53* copy number in elephants (20 copies) with several *TP53* retrogenes showing expression could be responsible for this phenotype [14, 15, 50]. We analyzed genomes of several bat species to see if *TP53* copy number expansion has occurred in Chiroptera. We found that several microbats have 2-4 copies of *TP53*. Most intriguingly, the vespertilionids in this study, *M. lucifugus,* and *E. fuscus,* seem to possess 7 and 2 copies of *TP53*, respectively. Of the seven copies of *TP53* in *M. lucifugus*, one is a full copy duplication and five are short retrocopies. Whether the p53 duplication plays role in enhanced basal p53 level remains to be determined. Elevated p53 activity is likely to contribute to cancer resistance in bats as increasing p53 dosage in mice significantly increased their resistance to cancer [62, 63].

In summary, our study demonstrates that bat fibroblasts undergo malignant transformation with just two oncogenic hits. Bat cells constitutively express telomerase and do not need to bypass replicative senescence barrier for malignant transformation. We also show that bats have elevated transcriptional levels and signaling through p53 pathways with some species showing genomic duplications (Figure 8). This adaptation may contribute to cancer resistance of bats but is unlikely to confer the same level of protection as achieved by extra 3 tumor suppressor barriers as observed in human. Bats have unique immune systems which allows them to survive a wide range of deadly viruses, and many unique immune adaptations have been described in bats (reviewed in [30]). We hypothesize, therefore, that bats rely more heavily on non-cell autonomous tumor suppressor mechanisms such as immune surveillance for elimination and detection of cancerous cells *in vivo*.

**Figure 8:**
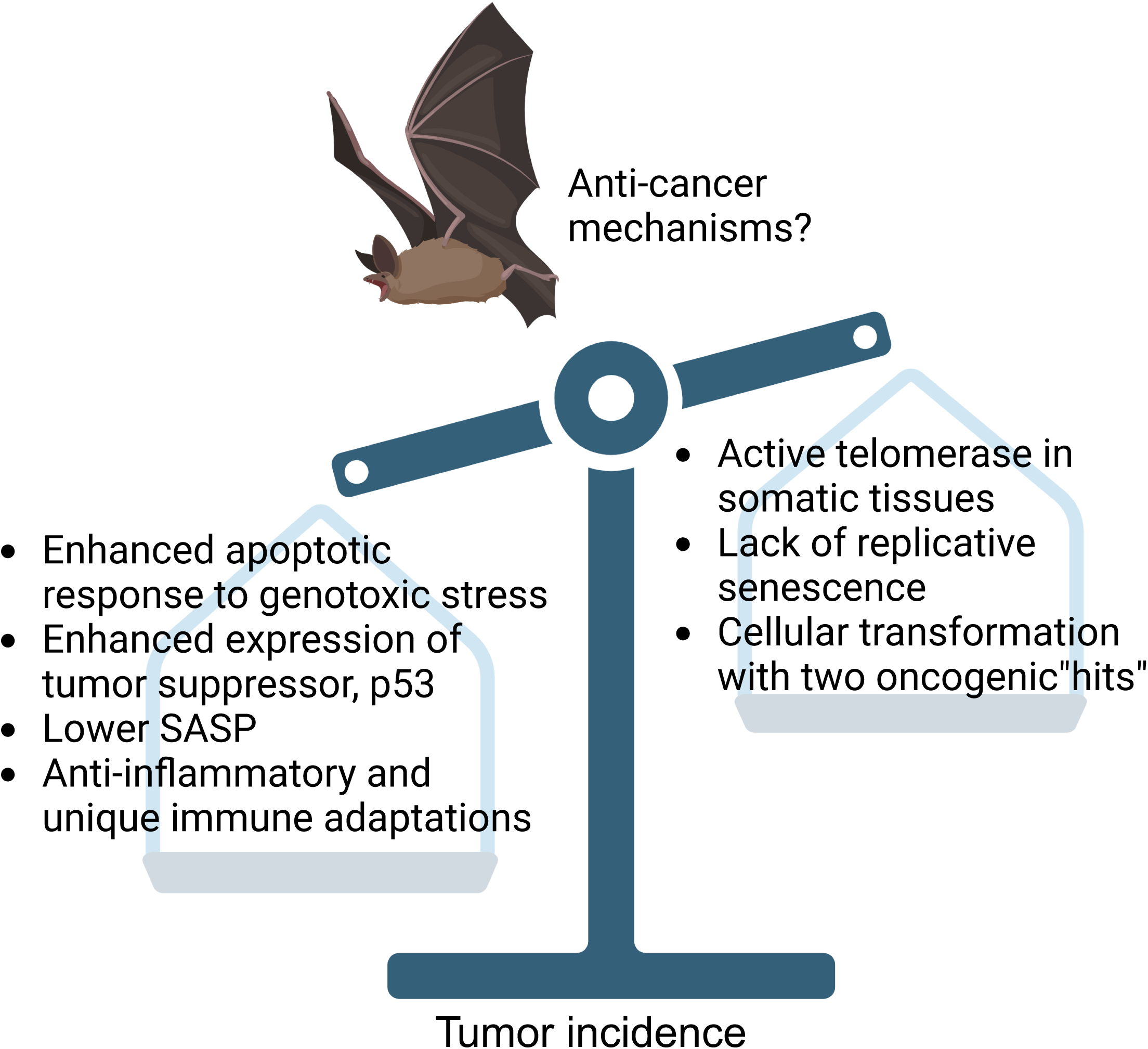
Graphical summary. Bat fibroblasts possess pro-tumorigenic characteristics like lack of replicative senescence, easy oncogenic transformation, and sustained telomerase activity. However, they also possess anti-tumorigenic properties like apoptotic response to genotoxic stress, lower SASP, and enhanced expression of tumor suppressor protein, p53. We hypothesize that a balance of these properties and non-cell autonomous factors like their unique immune system adaptations may contribute to their resistance to cancer.

## MATERIALS AND METHODS

### Animals

All animal experiments were approved by the University of Rochester Committee for Animal Research, Protocol number 2017-033.

*Myotis lucifugus* wing fibroblasts from three individuals were from Richard Miller collection at the University of Michigan. *Eptesicus fuscus* fibroblasts were isolated from wing tissue of four individual bats from the Northeast Ohio Medical University colony. *Eonycteris spelaea* fibroblasts were isolated from wing tissue of four individual bats from a breeding colony housed at Duke NUS in Singapore. *Artibeus jamaicensis* fibroblasts were isolated from wing tissue of four individual bats from the Colorado State University colony. Maximum lifespan (MLS) information for *M. lucifugus*, *E. fuscus*, and *A. jamaicensis* were obtained from AnAge [64]. MLS for *E. spelaea* is from personal communication from bat researchers. *Myotis myotis* were sampled in Morbihan, Brittany in North-West France, July 2021 in accordance with the permits and ethical guidelines issued by ‘Arrête ’ by the Préfet du Morbihan (Brittany) and the University College Dublin ethics committee. This population has been transponded and followed since 2010 as part of on-going mark-recapture studies by Bretagne Vivante and the Teeling laboratory.

### Cell culture

Primary fibroblasts were isolated from wing tissue of big brown bats, cave nectar bats, and Jamaican fruit bats, and little brown bats using the protocol described in detail previously [65]. *Myotis myotis* primary fibroblasts were generated and expanded from non-lethally sampled 3 mm wing biopsies using methods detailed in [1]. At least three individuals of each species were sampled generating independent cell lines. Isolated wing fibroblasts were grown in EMEM (ATCC) containing 15% fetal bovine serum (GIBCO), 100 U/mL penicillin, and 100 mg/mL streptomycin antibiotics (GIBCO). Culture conditions were 37°C with 5% CO_2_ and 5% O_2_. Cells were passaged when 80-90% confluent. Cells were trypsinized, and counted. Cells with PD lower than 30 were used for all experiments. Population doubling was calculated using the formula: [3.32*log (cell number harvested - cell number seeded)] + PD of cells seeded.

### Telomerase Repeated Amplification Protocol (TRAP)

TRAP assay was performed using TRAPeze kit (Cat. #S7700, Millipore Sigma). Briefly, 0.5 million cells were resuspended in CHAPs lysis buffer, and the amount of protein was estimated using Pierce BCA Protein Assay Kits (Cat. # 23225, Thermo Scientific). TRAP assay was performed using 250 ng of cell extracts, and manufacturer’s instructions were followed. HeLa cell lysates were used as positive controls.

### Soft Agar Assay

Following plasmids were used in this assay to generate stable cell lines: pCMV-HRas^G12V^ (Clontech), Piggyback vectors, pPB-SV40 LT, pPB-SV40 LT-K1, pPB-SV40 LT-Δ434-444, pBase, GFP control (Launchpad AVA2590, plasmid #85442). Cells were transfected using Amaxa nucleofector (Lonza) using the manufacturer’s protocol T-20. Stable cell lines were generated using puromycin (0.25-0.5 μg/mL) or hygromycin (25-50 μg/mL), depending on the plasmids and their combinations. For soft agar assay, 2X complete medium was prepared using 2X EMEM, 30% FBS, and 2X antibiotics as required. A base layer was poured into plates after mixing with 1% agarose (Difco Agar Noble). The top layer was made similarly, where 2X complete medium was mixed with 0.8% agarose and 10,000 cells/35 mm well and layered on top of the base layer. Following the solidification of agarose, 1 mL 1X complete media with antibiotics was added on top and refurbished twice a week. Plates were incubated at 37°C in a humidified incubator for three weeks. Plates were then imaged for colonies in soft agar.

### Xenograft assay

Animal experiments were performed under pre-approved protocols and in accordance with guidelines set by the University of Rochester Committee on Animal Resources (UCAR). Fibroblasts stably expressing oncogenes or a combination of oncogenes were tested for tumor formation in nude mice (Charles River, Crl :NIH-Lystbg-JFoxn1nuBtkxid). Each flank of the nude mice was injected with two million cells in 100 μL PBS mixed with an equal volume of Matrigel using a 22-gauge needle. A total of 8 injections per cell line were tested. Tumor formation was monitored twice a week, and dimensions were measured using Vernier calipers. Tumor long diameter < 5 mm was considered negative, and 20 mm was considered the tumor burden endpoint. Mice that did not reach tumor burden endpoints were terminated after a maximum of 60 days. Euthanized mice were photographed, and tumors were excised, photographed, weighed, frozen at -80°C, and preserved in formalin.

### Gamma-Radiation and Nutlin-3 treatments

Radiation treatment was performed using γ-radiator (Model 8114 Shepherd Cs^137^). Cells were seeded 24-48 h before treatments. Following treatment with radiation doses 10 and 20 Gy, cell culture medium was immediately changed, and cells were returned to the incubator.

For induction of senescence, radiated cells were incubated for 12 days, and the culture medium was replaced twice a week. For Nutlin-3 (CAS 548472-68-0, Santa Cruz) treatment, different concentrations of Nutlin-3 (10, and 50 μM) in DMSO were added to cells and incubated for 24 h for apoptosis assay or 6 h for western blotting.

### BrdU incorporation assay

Cells were seeded 24 h before treatment and were allowed to grow in the presence of 3 μg/mL BrdU (BD Pharmingen) for 48 h. Cells were trypsinized and fixed with 70% cold ethanol for 30 min. For staining, fixed cells were washed twice with PBS and treated with 2 N HCl for 30 min for DNA denaturation. Following two more PBS washes, cells were incubated with 5% BSA for 1 h at room temperature (RT). Cells were then incubated with anti-BrdU (Alexa Fluor® 647 Mouse anti-BrdU, Clone 3D4 (RUO)) antibody overnight at 4°C. Following 2x PBS washes, stained cells were analyzed by flow cytometry with appropriate positive and negative controls.

### Apoptosis assay

Apoptosis assay was performed using Annexin V FLUOS staining kit (Roche), and the manufacturer’s instructions were followed with slight modifications. Briefly, culture supernatant and PBS washes were collected. Cells were then trypsinized briefly, and the cell pellet was mixed with the floating cells collected from the supernatant and washes. Following two more washes with PBS, the cell pellet was resuspended in Annexin-V-FLUOS labeling reagent containing Annexin V and propidium iodide. Following a 10 min incubation on ice, cells were immediately analyzed by flow cytometry with appropriate positive and negative controls.

### Senescence-associated **β**-gal staining

Irradiated cells were incubated for 12 days, and the media was replaced twice a week. After 12 days, approximately 24-48 h before staining, cells were seeded to be approximately 40% confluent. Cells were washed 2X with PBS and fixed for 5 min with 2% formaldehyde. Cells were washed gently to remove formaldehyde and staining solution (1 mg/ml X-gal in DMSO, 40 mM Citric acid/Sodium phosphate buffer pH 6, 5 mM potassium ferricyanide, 5 mM potassium ferricyanide, 150 mM NaCl, 2 mM MgCl_2_) was added to cells. Cells were incubated for 12-24 h in a 37°C. Following the development of a blue color, cells were washed 2X with PBS, overlayed with 70% glycerol, and imaged. Cells were counted under light microscope, and the percentage of SA-β-gal positive cells was determined.

### Plasmid luciferase reporter assay

Cells were seeded in 24-well plates 24 h before transfection. Following manufacturers’ instructions, about 250-300 ng of plasmids were transfected using Fugene (Invitrogen). p53-TA-Luc (Clontech/Takara) was used to assay p53 transcriptional activity and pRL-CMV (Promega) was used as the transfection control. Reporter activity was analyzed 48 h post-transfection using the Dual luciferase reporter assay system and GloMax 20/20 Luminometer (Promega) instrument. Manufacturers’ instructions were followed, and activity ratios were reported as relative luciferase units (RLU).

### RNA sequencing (RNA-seq)

Total RNA was extracted from irradiated or non-irradiated fibroblasts from four bat species, laboratory mice, wild-caught mice and humans using PureLink^TM^ RNA Mini Kit (Thermo Fisher Scientific) following manufacturer’s instructions. NGS-TruSeq Stranded mRNA libraries were generated and sequenced with Illumina NovaSeq 6000 single-end 75 bp sequencing at the University of Rochester Genomics Research Center.

### Genome assembly, annotation, and gene expression analysis

Raw reads were demultiplexed using configurebcl2fastq.pl (v1.8.4). Adapter sequences and low-quality base calls (threshold: Phred quality score <20) in the RNA-seq reads were first trimmed using Fastp (0.23.4)[66]. For all species, the clean reads were aligned using Salmon (v1.5.1) to the longest coding sequence (CDS) of each gene extracted from corresponding genome assembly based on genome annotations using GffRead [67]. The genome assemblies used in this study include GCA_003508835.1 (*Eonycteris spelaea*), EptFus1.0_HiC (DNA Zoo Consortium, *Eptesicus fuscus*), GCA_004027435.1 (*Artibeus jamaicensis*), Myoluc2.0_HiC (DNA Zoo Consortium, *Myotis lucifugus*), GCF_000001635.27 (*Mus musculus*), and GCA_000001405.29 (*Homo sapiens*). Human-referenced TOGA annotations were used when the original genomic annotations are not available [68]. The orthologous genes of each species to human reference were identified by performing reciprocal blast search (BLAST+ v2.10.1) [69] against human longest protein (GRCh38.p13; Ensembl database, release109) with parameters of “-evalue 1e-05; -max_target_seqs 1”, and hits with query coverage > 30% were retained. The values of read count and effective gene lengths for each gene were collected and integrated into gene-sample table according to their orthologous relationship. Salmon transcript counts were used to perform differential expression analysis. Only human genes with orthologs in all species were kept for the downstream analysis. To filter out low expressed genes, only genes with all sample read counts sum >10 were retained. The filtered count matrix was normalized using median of ratios method [70] implemented in DESeq2 package [71]. Since orthologous gene lengths could vary among different species, we implemented an additional length normalization step in the DESeq2 pipeline to avoid biased comparative quantifications resulting from species-specific transcript length variations. To do this, the matrix of effective lengths for each gene in each sample was delivered to the DESeq2 ‘DESeqDataSet’ object so that they are included in the normalization for downstream analysis. Differential expression analysis, using either irradiated cells (24 h or 12 days) versus control (24 h or 12 days) or only irradiated cells from 24 h versus 12 days in each species, was performed using DESeq2. We considered all differentially expressed genes (DEGs) with an adjusted p < 0.05, and 1.5-fold expression change to be statistically significant in our analyses.

### Functional enrichment analysis

Hallmark gene sets and C2 curated gene sets from MsigDB were used for GSEA analysis using the ranked fold changes from the DEG analysis. The *p*-values were calculated using a permutation-based approach. For cellular senescence analysis, in addition to MsigDB gene sets, we also collected gene sets identified as the senescence signature using transcriptomic and proteomic approaches from published literature [40–42].

### TP53 copy number analysis in *M. lucifugus* genome

BLAST was used to search for *TP53* genes in bats genomes using the human TP53 protein sequences as a query. The best matched genomic regions plus 5kb up- and downstream flanking sequence were extracted from the genome for GeneWise gene structure prediction. Predicted genes with more than 80% converge of human TP53 protein were treated as the *TP53* duplicates. To confirm that the identified TP53 copies in *M. lucifugus* are not misidentified homologs from the p53 gene family, additional blast analysis with human *TP63* and *TP73* was performed.

### Cas9 targeted sequencing

High molecular weight (HMW) genomic DNA from a single *M. lucifugus* fibroblast line was isolated using the Nanobind ®HMW DNA extraction kit for cultured cells (PacBio). *M. lucifugus* TP53 copy sequences found in the Myoluc2.0 genome assembly were used to design guide RNAs (listed in Supplementary Table 2) using Benchling. Guides were designed to cut within sequences of TP53 gene copies and sequence across part of the gene into flanking regions. The Cas9 Sequencing Kit SQK-CS9109 (Oxford Nanopore Technologies) was used for targeted sequencing according to manufacturer protocol. Briefly, extracted HMW gDNA was phosphatase treated to remove pre-existing phosphorylated ends, followed by heat inactivation. Next, target regions were cleaved in vitro using Cas9 ribonucleoprotein complexes consisting of Alt-R® S.p. HiFi Cas9 Nuclease V3 (Integrated DNA Technologies), Alt-R® CRISPR-Cas9 tracRNA (Integrated DNA Technologies), and target-specific Alt-R® CRISPR-Cas9 crRNAs (Integrated DNA Technologies). Following cleavage, Cas9 was heat inactivated and adapters were ligated to cleaved phosphorylated ends. Adapters from Ligation Sequencing Kit V14 SQK-LSK114 (Oxford Nanopore Technologies) were used in place of the adapter included in the Cas9 kit. Following adapter ligation, samples were cleaned up using AMPure XP Beads and loaded onto a R10.4.1 flow cell on a MinION Mk1C (Oxford Nanopore Technologies) for sequencing. Raw data was basecalled in Super-High accuracy mode and aligned to *M. lucifugus* TP53 reference sequences in both Guppy and Dorado with read splitting enabled. List of guides used is shown in Supplementary table 2.

## Acknowledgements

We thank Randy Foo for providing technical assistance with bat colony management at Duke-NUS and providing bat tissues for this study. The work at Duke-NUS is funded by grants from Singapore National Research Foundation (NRF-CRP10-2012-05), National Medical Research Council (OFIRG19nov-0050) and Ministry of Education (MOE2019-T2-2-130). This study was supported by grants from the National Institute on Aging to VG and AS, and grants from Michael Antonov Foundation and Milky Way Research Foundation to VG and from a Science Foundation Future Frontiers (grant no. 19/FFP/6790) awarded to ECT.

## Supplementary figures

**Supplementary Figure 1: PCA and Pearson correlation matrix for samples from all species, and expression of genes regulating TP53**

a,b, PCA plots and Hierarchical clustering analysis showing that samples cluster by species, but not treatment and timepoint. c,d Boxplots showing the expression changes of (c), MDM2 and (d) WRAP53 in each species day 12 post-radiation. The box plots display the median, the1st, and 3rd quartiles; the whiskers show a1.5×interquartile range. Data points outside the whiskers are outliers. e, TRAP assay showing telomerase activity in *Myotis myotis* wing fibroblasts.

**Supplementary Figure 2: Expression of SASP genes in bats when compared to mice**

Scatter plots showing the expression differences between 4 bat species and mouse for genes in 7 SASP-related gene sets 12 days after irradiation (see Methods for gene sets). Percentages in left-upper and right-lower side indicates the fraction of genes with higher expression in mouse and *M. lucifugus*, respectively.

**Supplementary Table 1: Details of bats used in the study**

Details of bats used in the study showing characteristics of each bat.

**Supplementary Table 2: Guide RNAs used in targeted long-read sequencing of p53 in *M. lucifugus***

Sequences of targeted guide RNAs used in Cas9-mediated targeted sequencing of TP53 sequence long reads in *M. lucifugus*.

