## Supplementary Figure 1 for "Limited Cell-Autonomous Anticancer Mechanisms in Long-Lived Bats"

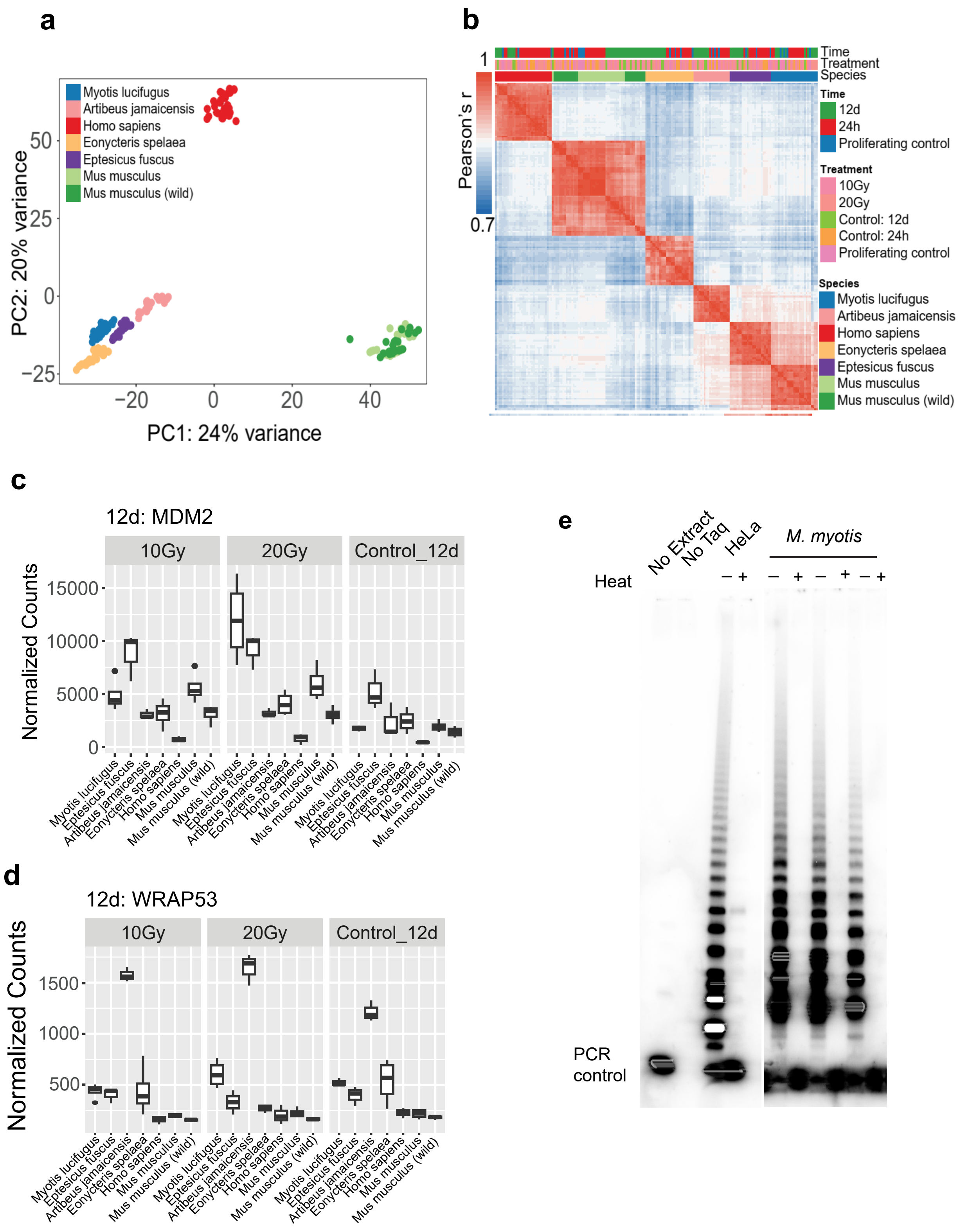


**Supplementary Figure 1: PCA and Pearson correlation matrix for samples from all species, and expression of genes regulating TP53**

a,b, PCA plots and Hierarchical clustering analysis showing that samples cluster by species, but not treatment and timepoint. c,d Boxplots showing the expression changes of (c), MDM2 and (d) WRAP53 in each species day 12 post-radiation. The box plots display the median, the1st, and 3rd quartiles; the whiskers show a1.5×interquartile range. Data points outside the whiskers are outliers. e, TRAP assay showing telomerase activity in *Myotis myotis* wing fibroblasts.
