## Supplementary Figure 2 for "Limited Cell-Autonomous Anticancer Mechanisms in Long-Lived Bats"

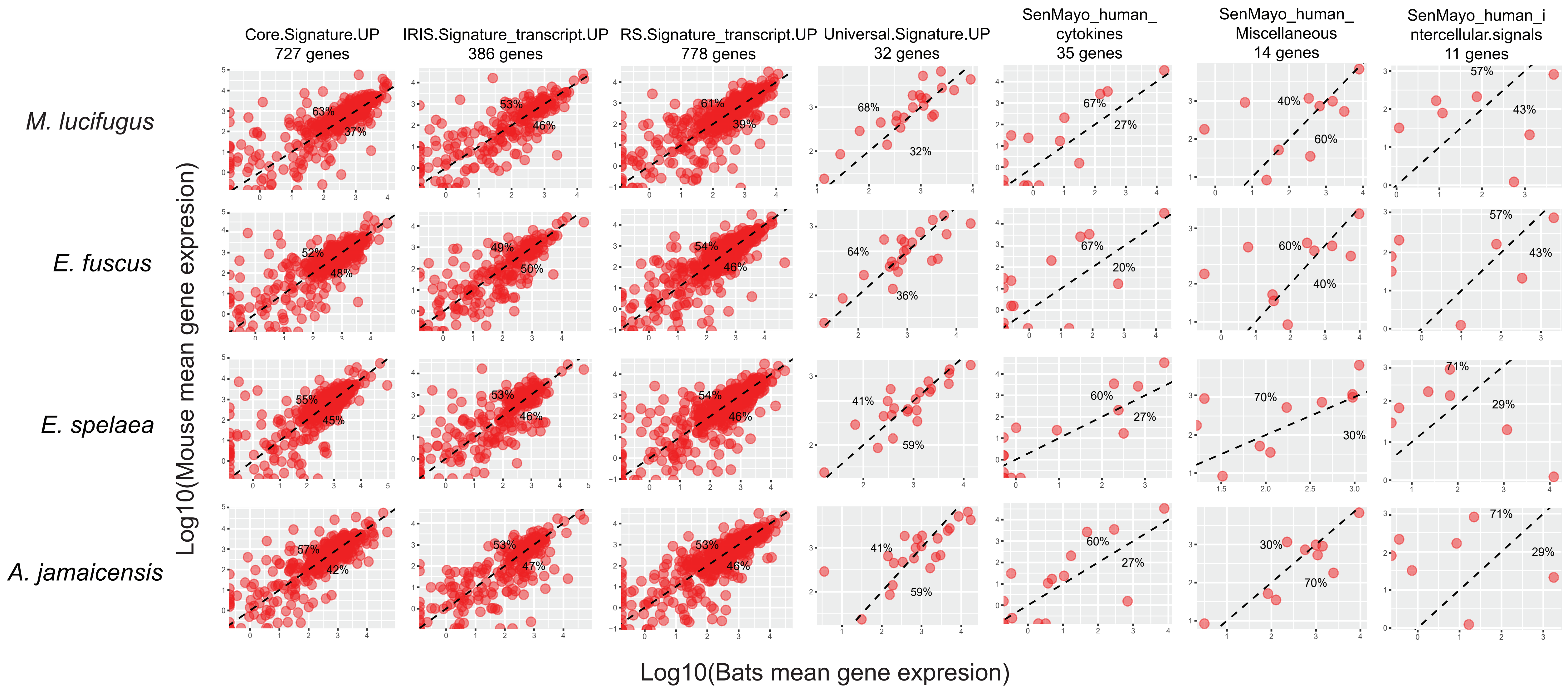


**Supplementary Figure 2: Expression of SASP genes in bats when compared to mice**

Scatter plots showing the expression differences between 4 bat species and mouse for genes in 7 SASP-related gene sets 12 days after irradiation (see Methods for gene sets). Percentages in left-upper and right-lower side indicates the fraction of genes with higher expression in mouse and M. lucifugus, respectively.
