## Supplementary Table 1 for "Limited Cell-Autonomous Anticancer Mechanisms in Long-Lived Bats"

| Common name | Little brown bat | Big brown bat | Cave nectar bat | Jamaican fruit bat |
|---|---|---|---|---|
| Species name | *Myotis lucifugus* (*M. lucifugus*) | *Epstesicus fuscus* (*E. fuscus*) | *Eonycteris spelaea* (*E. spelaea*) | *Artibeus jamaicensis* (*A. jamaicensis*) |
| MLS (wild) | 34 years | 19 years | approx. 10 years (captive) | 19.2 years |
| Body temperature | 32° C | 36° C | 34° C | 36.4 |
| Adult weight | 10 g | 23 g | Upto 60 g | 42 g |
| Hibernation | yes | yes | no | no |
| Diet | insects | Animal, insects | Fruit, nectar | Fruit |
| Habitat type | throughout North America | throughout North America | South Asia, tropical | Southern Mexico, South America, Caribbean Islands, Southern Bahamas |
| Progeny/year | 1 | 2 | 2 | 1 |
| Cave roosting | Sometimes | Sometimes | Yes, exclusively | Yes |

**Supplementary Table 1: Details of bats used in the study**

Details of bats used in the study showing characteristics of each bat
