## Supplementary Table 2 for "Limited Cell-Autonomous Anticancer Mechanisms in Long-Lived Bats"

| Guide RNA | Sequence |
|---|---|
| In-F-canon-1 | GCC CCU UUU UAG GUU UCA GGG |
| In-F-canon-2 | CCA UCA GAG UGA GAG GUG UAG |
| In-R-canon-1 | GAG GUC UGG UUU GGC AAC UGG |
| In-R-canon-2 | CAC CCU CCC CUC ACA GUA CGG |
| In-F-retro-1 | GGA GUC UUU CAG CAU GAU GAG |
| In-F-retro-1a | GGA AUC UUC CAG UGU GAU GAG |
| In-F-retro-2 | GCU CAA UGA AUG CUC CCA GUG |
| In-F-retro-2a | GCU UGC UUA GUG CUU CCA GGG |
| In-F-retro-2b | GUU CAA UUC AUG CUC CCA GUG |
| In-F-retro-3 | UGC UCA AUG AAU GCU CCC AGG |
| In-F-retro-3a | UGC UUG CUU AGU GCU UCC AGG |
| In-R-retro-1 | AAG ACC UAC CCU GAC AGC UAG |
| In-R-retro-1a | AAG ACC UAU CCU GGU AGC UAG |
| In-R-retro-2 | GAC AGC UAU GGU GUC UGU UUG |
| In-R-retro-2a | GGU AGC UAU GGU UUC CGU UUG |
| Pair-F1 | GAA UGU AUG GAG UUG UAA UCG |
| Pair-F2 | GCU AUA UAC CAA UCU GCA GAG |
| Pair-F3 | AAG GGU GGC GGU GUC AAC AGG |
| Pair-R1 | UCC GGU CAU GAC ACA UAC UGG |
| Pair-R2 | AUU CCG GUC AUG ACA CAU ACG |
| Pair-R3 | GGG UUG UGG AUU CGA UUC AGG |

**Supplementary Table 2: Guide RNAs used in targeted long-read sequencing of p53 in *M. lucifugus***

Sequences of targeted guide RNAs used in Cas9-mediated targeted sequencing of TP53 sequence long reads in *M. lucifugus*.
